# Cold survival and its molecular mechanisms in a locally adapted nematode population

**DOI:** 10.1101/2021.05.10.443188

**Authors:** Wenke Wang, Anna G. Flury, Jennifer L. Garrison, Rachel B. Brem

## Abstract

Since Darwin, evolutionary biologists have sought to understand the drivers and mechanisms of natural trait diversity. The field advances toward this goal with the discovery of phenotypes that vary in the wild, their relationship to ecology, and their underlying genes. Here, we established resistance to extreme low temperature in the free-living nematode *Caenorhabditis briggsae* as an ecological and evolutionary model system. We found that *C. briggsae* strains of temperate origin were strikingly more cold-resistant than those isolated from tropical localities. Transcriptional profiling revealed expression patterns unique to the resistant temperate ecotype, including dozens of genes expressed at high levels even after multiple days of cold-induced physiological slowdown. Mutational analysis validated a role in cold resistance for seven such genes. As the temperate *C. briggsae* population likely diverged only ~700 years ago from tropical ancestors, our findings highlight a candidate case of very rapid, robust, and genetically complex adaptation, and shed light on the mechanisms at play.

## Introduction

Understanding how and why organisms differ in the wild is a key goal of evolutionary biology. Some traits are evolutionary accidents, and others arise under local adaptation, fixing in a population when they promote fitness in a new niche. Dissecting these processes requires case studies where we can establish the underlying ecology, the phenotypes that have evolved, and, ultimately, the molecular mechanisms. Invertebrates can provide exceptional power toward this end (Adrion, Hahn, & Cooper, 2015; Kraemer & Boynton, 2017; Sanford & Kelly, 2011; Savolainen, Lascoux, & Merilä, 2013), but even in lower eukaryotes, genetically tractable ecological study systems remain at a premium in the field. Of particular interest are cases in which tools from the lab can be brought to bear to study evolution in natural settings.

*Caenorhabditis briggsae,* like its relative the model organism *C. elegans*, is a free-living nematode that has been isolated all over the world (Baird & Stonesifer, 2012; Cutter, Félix, Barrière, & Charlesworth, 2006; Okahata et al., 2016; Prasad, Croydon-Sugarman, Murray, & Cutter, 2011; Stegeman, De Mesquita, Ryu, & Cutter, 2013). Genetic and phenotypic analyses have revealed a split between strains of *C. briggsae* isolated from temperate and tropical localities (Cutter et al., 2006; Graustein, Caspar, Walters, & Palopoli, 2002). The contrast between these populations serves as a useful framework for the study of ecological diversification. Under current models, ancestral *C. briggsae* occupied the tropical niche, with colonization of and recent expansion into temperate latitudes ~700 years ago, possibly associated with human activity (Cutter et al., 2006). Elegant reports have characterized differences between temperate and tropical *C. briggsae* in temperature-dependent fecundity (Prasad et al., 2011) and behavior (Baird & Stonesifer, 2012; Cutter et al., 2006; Stegeman, Baird, Ryu, & Cutter, 2019; Stegeman et al., 2013), with a focus on chronic response to warm and hot conditions (14°C to 30°C). Resistance to acute temperature shock, as it has evolved among *C. briggsae* populations, remains less well understood. The response to extreme low temperature is a compelling potential character under ecological pressures in the *C. briggsae* system, as it is likely to manifest in the winter season of temperate latitudes and not in tropical regions (Lacher & Goldstein, 1997).

In this study, we investigated the response to cold stress across *C. briggsae* strains isolated from different niches. We found that temperate strains survived cold conditions in which most animals of tropical origin died. Using two strains, AF16 and HK104, as representative of the tropical and temperate populations respectively, we profiled cold-responsive transcriptomes to achieve molecular insight into the divergence in the cold resistance trait. Against a backdrop of thousands of genes with cold-evoked expression patterns shared between the strains, we found >100 genes with high expression unique to cold-treated HK104. In mutational tests, we confirmed the role of seven of the latter genes in cold resistance.

## Materials and methods

### Worm strains

Wild-type strains of *C. briggsae* (AF16, VT847, ED3083, JU726, QX1410, HK104, PB826, JU439, EG4181 and VX34) and *C. elegans* (CB4555, N2, AB1, JU262, PX179, JU258, JU1172, JU1652, JU393, MY16, JU779, JU1088, GXW1, ED3077 and ED3052) are described in Table 1 and Supplementary Table 2 respectively. To generate and transcriptionally profile AF16 x HK104 F1 hybrids, we first made a marked strain of AF16, CP161 (*Cbr-unc-119(nm67) III; nmIs7 [Cni-mss-1(+) + Cni-mss-2(+) + Cbr-myo-2::GFP + unc-119(+)]*). We then crossed HK104 hermaphrodites with CP161 males, picked labeled hermaphrodite F1 progeny at the L4 stage, and used them as input into cold treatment and expression profiling procedures as detailed below.

**Table 1.** Collection localities of tropical and temperate C. briggsae strains.

| Strain | Locality of origin | Phylogeographic group | Latitude | Approximate elevation (m) | Mean winter temperature (°C min, max) | Mean summer temperature (°C min, max) |
| --- | --- | --- | --- | --- | --- | --- |
| AF16 | Ahmedabad, India | Tropical | 23° 01'N | 50 | 14.4, 29.4 | 26.1, 33.9 |
| VT847 | Hawaii, USA | Tropical | 20° 57'N | 930 | 18.9, 26.1 | 22.8, 29.4 |
| ED3083 | Johannesburg, S. Africa | Tropical | 26° 10'N | 1,750 | 2.8, 18.9 | 15.6, 27.2 |
| JU726 | Chengyang Village, China | Tropical | 44° 28'N | 11 | 3.9, 12.8 | 24.4, 34.4 |
| QX1410 | St. Lucia | Tropical | 13° 22'N | 14 | 23.0, 27.0 | 25.0, 31.0 |
| HK104 | Okayama, Japan | Temperate | 34° 40'N | 30 | 0, 8.3 | 22.8, 30.6 |
| PB826 | Ohio, USA | Temperate | 39° 34'N | 270 | -6.1, 2.8 | 17.8, 30 |
| JU439 | Reykjavik, Iceland | Temperate | 64° 08'N | 16 | -2.2, 2.2 | 8.9, 14.4 |
| EG4181 | Salt Lake City, Utah, USA | Temperate | 40° 42'N | 1,290 | -7.8, 2.8 | 15.6, 34.4 |
| VX34 | Guangshui, Hubei, China | Temperate | 30° 58'N | 80 | -6.1, 0 | 13.8, 23.9 |

*C. elegans* mutant strains profiled in Figure 5 are as follows: JT366 *vhp-1(sa366) II*, IG685 *tir-1(tm3036) III*, KU4 *sek-1(km4) X*, NL152 *pgp-1(pk17) IV*; *pgp-3(pk18) X*; *mrp-1(pk89) X*, RB1916 *pgp-8(ok2489) X*, RB1840 *M28.8(ok2380) II*, VC2677 *Y47G6A.5(gk1098) I*, GH403 *glo-3(kx94) X*, VC422 *tag-120(gk221) V*, VC1392 *zip-5(gk646) V*, RB792 *F09C12.2(ok582) II*, RB1284 *C30F12.6(ok1381) I*, VC2072 *grh-1(gk960) I*, RB2200 *gst-24(ok2980) II*, VC1499 *nhr-117(gk707) V*, JT5244 *aex-4(sa22) X*, RB1267 *D1009.3(ok1349) X*, RB2171 *Y4C6A.1(ok2938) IV*, VC2214 *nhr-178(gk1005) V*, RB1749 *numr-1(ok2239) III*, RB2499 *T27F2.4(ok3462) V*, RB1067 *his-24(ok1024) X*, VC1544 *C12D12.5(gk700) X*, IG544 *nipi-3(fr4) X*, RB1362 *H22K11.4(ok1529) X*, BC15170 *dpy-5(e907) I*, COP677 *ncr-1(knu4) X*, *swt-6(tm5930) V*, *K03H1.5(tm10908) III*, *lips-10(tm7601) II*, *F26A10.2(tm549) X*, *F45E10.2(tm5965) II*, *arrd-25(tm12435) V*, *C18H9.5;C18H9.1(tm12783) II*, *bigr-1(tm6317) II*, *smoc-1(tm7000) V*, *npr-13(tm1504) V*, *sek-3(tm1344) X*, *cnp-3(tm2950) X*, *fpn-1.1*; *npp-13(tm6914) I*, *osta-3(tm5747) II*, *C18H7.11(tm12727) IV*, *F21G4.1; mrp-4(tm10068) X*.

### Cold tolerance assays

Cold tolerance assays were performed as described (Jiang et al., 2018). For a given biological replicate of a given strain or treatment condition, 20-30 worms were dispersed on each of 2-3 plates and raised at 20°C until 24 hours after they reached the L4 stage. Plates were then distributed with equal distance between them in a box and transferred to a constant 4°C cold room for the duration indicated below and in figure captions.

Plates were then moved to room temperature for 2 hours before survival scoring, in which worms that failed to respond to a gentle prodding with a platinum wire were scored as dead. For each strain at least three independent experiments were performed.

### RNA isolation and library preparation for RNA-seq

RNA isolation was performed essentially as described (Wang et al., 2018). For a given biological replicate of a given strain, 100-200 synchronized mid/late L4 stage worms were picked and incubated at 20°C for 24 hours, after which one cohort of worms was harvested immediately, representing the untreated control, and another cohort was subjected to 4°C treatment for 60 hours as above, followed by harvest. Two replicates for each strain and each condition were performed. Collected worms were homogenized in 1 ml Tri-reagent (ThermoFisher) for 30 min at room temperature. 0.1 mL of bromochloropropane (BCP, Sigma) was added to the sample and mixed well. The sample was then spun at 12,000g for 15 min at 4°C and the aqueous phase was transferred to a new tube. 0.5 mL Isopropanol was added to the sample, followed by incubation at room temperature for 10 min and centrifugation at 12,000g for 10 min. The RNA pellet was washed twice with 75% EtOH and dissolved in water. The RNA sample was then purified to remove DNA using the RNeasy Mini Kit (Qiagen), and used as input into RNA-seq library preparation using KAPA RNA hyper prep kit (Roche).

### Amended reference genome construction

We established a reference genome for *C. briggsae* strain HK104 as follows. Raw genome sequencing reads for HK104 were downloaded from the NCBI (project accession PRJNA509247). These reads were aligned to genome assembly CB4 of the reference sequence of the *C. briggsae* AF16 strain (Genbank Assembly Accession GCA_000004555.3) using bowtie2 with default parameters. Samtools, bcftools, and bgzip were used to call SNPs, retaining those with a quality score of >20 and combined depth of >5 and <71. We then generated a pseudogenome by replacing the reference AF16 allele with that of HK104 at each SNP using bcftools, totaling 441,227 SNPs. This new amended HK104 genome was then concatenated to the reference AF16 genome to form a master AF16-HK104 genome.

### RNA-seq data analysis

RNA-seq data analysis was essentially as described (Wang et al., 2018). For each library, ~20M 150bp length paired-end reads were generated. Low quality reads were removed using the FASTX Toolkit. Illumina primer sequences (adaptors) were removed from read sequences using cutadapt.

For transcriptomes of purebred AF16 and HK104, the resulting trimmed reads were aligned to the respective reference genome using the HISAT2 2.2.0 alignment program with default parameters. Mapped reads were then input into HTSeq 0.11.1 to calculate normalized counts for each gene. Genes with fewer than 20 reads mapped to them in all samples were removed and the remaining genes were used as input to test for differential expression using the generalized linear model framework in edgeR (in the module glmQLFTest) as follows. For Figure 3, each strain’s transcriptomes were used as input to a test for genes whose expression changed between cold and untreated samples. For Figure 4, the complete set of transcriptomes from both strains and both conditions was used as input to a test for genes at which the impact of cold on expression was different between the strains.

For transcriptomes of the AF16 x HK104 F1 hybrid, trimmed reads were aligned to a concatenation of the two strains’ reference genomes, and only reads that mapped to a single location in this concatenated reference were retained for analysis, reflecting allele-specific expression at the respective strain’s allele of the respective gene. Normalized read counts and significance testing were as for Figure 4 above, reporting cases of temperature effects that differed between the two strains’ alleles.

All raw transcriptome data are available at GEO: GSE171725 and SRA: SRP314054.

Principal component analysis was performed using the built-in function in edgeR. Gene Ontology term enrichment analysis was performed using the web tool at geneontology.org.

Genes for the mutant screen of Figure 5 in *C. elegans* were chosen as those that exhibited higher expression under cold treatment than in control condition in HK104 when analyzed separately, at nominal *p* < 0.01; exhibited stronger cold-evoked expression change in HK104 than in AF16 in the strain-by-temperature analysis, at nominal *p* < 0.01; and were annotated with one-to-one orthology between *C. elegans* and *C. briggsae*.

### CRISPR-mediated genome editing

CRISPR-mediated genome editing for *C. briggsae* was performed using an established protocol (Culp et al., 2020) modified from the analogous protocol for *C. elegans* (Friedland et al., 2013). For generating mutations in WBGene00025434, WBGene00037162, WBGene00025987, WBGene00034955, and WBGene00031437, the *C. briggsae* orthologs of *vhe-1*, *pgp-8, M28.8, ncr-1 and K03H1.5* genes respectively, day 1 adult animals of AF16 and HK104 were injected with pDD162 (*Peft-3::Cas9::tbb-2 3’UTR*), pCFJ90 (*Pmyo-2::mCherry*) and 2-4 different pU6::sgRNAs for the gene of interest. Surviving worms were separated, and F1 mCherry-positive animals were collected; their progeny, representing the F2 generation, were genotyped for the respective gene. Mutant genotypes are reported in Supplementary Table 3.

## Results

### Wild C. briggsae isolated from temperate but not tropical regions survive hypothermia independent of rearing temperature

Since temperature is one of the major factors that distinguish temperate and tropical climates (Lacher & Goldstein, 1997; Prasad et al., 2011), we hypothesized that *C. briggsae* strains from these two niches would respond to hypothermia differently. To test this, we cultivated *C. briggsae* strains from the tropical cluster (AF16, VT847, ED3083, JU726 and QX1410) and strains from the temperate cluster (HK104, PB826, JU439, EG4181 and VX34) (Table 1) at 20°C until they reached adulthood, and then switched the animals to 4°C. All worms were immobile after 60 hours of cold treatment, but upon recovery, strains from the temperate cluster survived at ~100%, whereas strains from the tropical cluster died at high rates (Figure 1).

**Figure 1.**
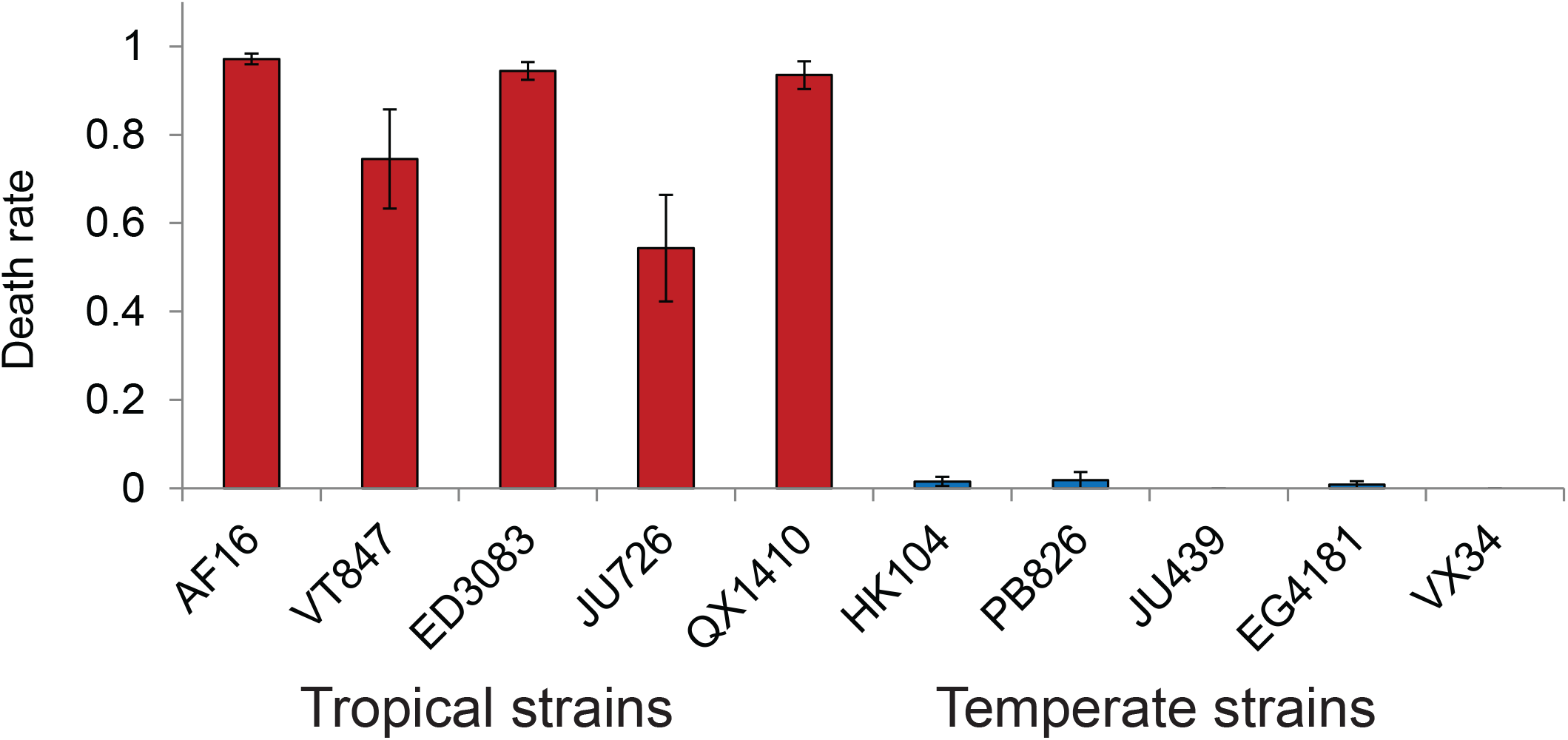
Cold survival differentiates tropical and temperate *C. briggsae*. Each bar reports the mean proportion of animals of the indicated *C. briggsae* strain surviving after development at 20°C followed by 60 hours of incubation at 4°C, across ≥ 3 biological replicates. Error bars indicate ±1 standard error of the mean. See Table 1 for strain information.

Cultivation temperature during development can affect the cold tolerance of adult worms (Ohta, Ujisawa, Sonoda, & Kuhara, 2014; Okahata et al., 2016). To examine this effect in the *C. briggsae* system, we used AF16, isolated from tropical India, and HK104, from a temperate niche in Japan, as representatives of the respective clades. As expected (Ohta et al., 2014), cold tolerance decreased in both AF16 and HK104 animals that had gone through development at higher temperatures (Figure 2) and in worms treated with longer cold shock (Supplementary Figure 1). However, at all rearing temperatures, HK104 was far more likely to survive cold treatment than was AF16 (Figure 2). Together, these results reveal protection from lethal cold shock as a trait specific to temperate-clade *C. briggsae*, in a manner that is largely independent of rearing temperature.

**Figure 2.**
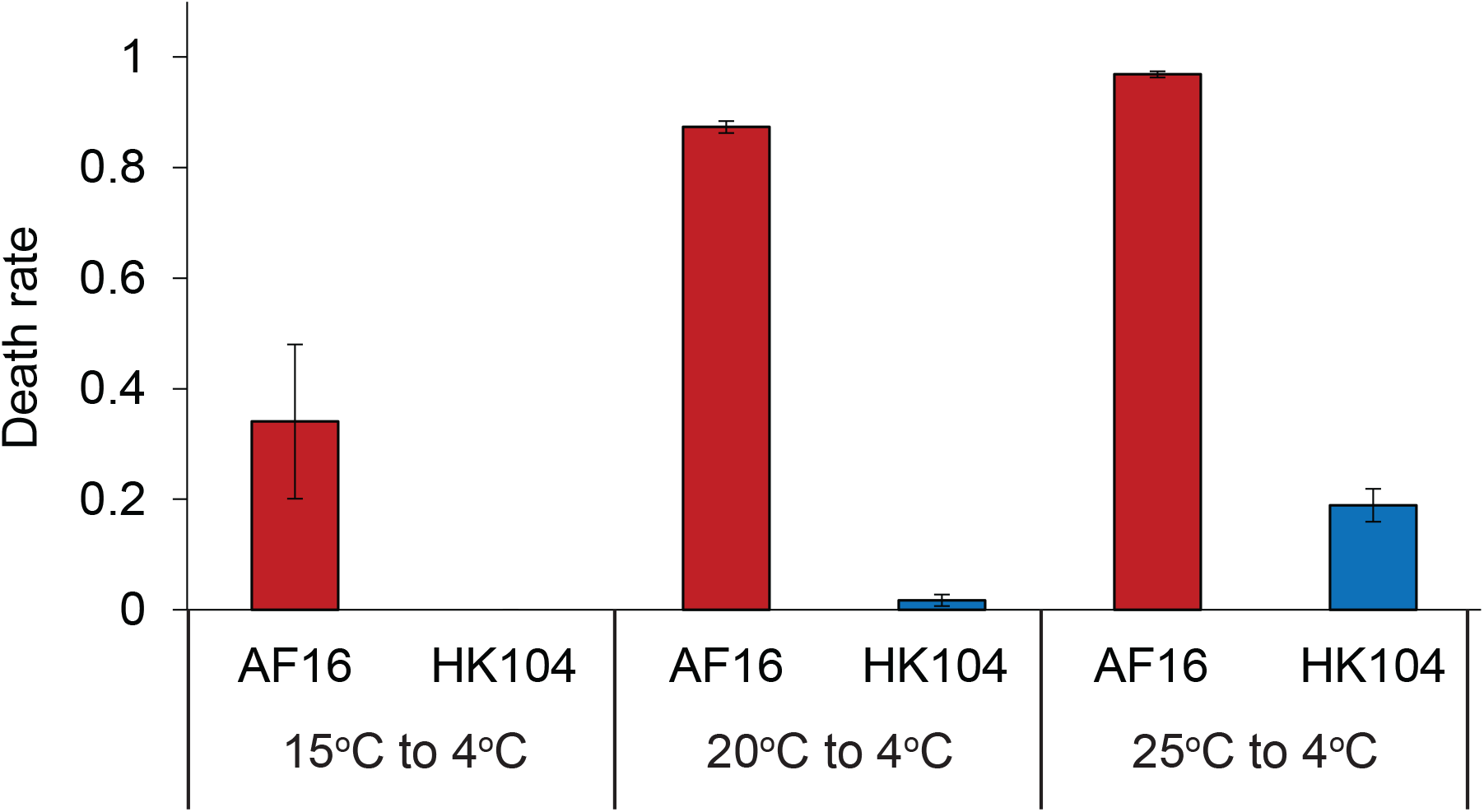
Temperate *C. briggsae* strain HK104 is much more cold-tolerant than tropical strain AF16 regardless of rearing temperature. Symbols are as in Figure 1, except that rearing temperature was 15°C (left), 20°C (middle), or 25°C (right) before cold shock.

### Transcriptional responses to cold stress unique to and shared between *C. briggsae* strains

To gain molecular insight into differences in cold stress response between tropical- and temperate-clade *C. briggsae*, we took a transcriptional approach, again making use of the AF16 and HK104 isolates as a model comparison. Transcriptome profiles of these strains before and after cold treatment revealed robust clustering by temperature and strain background, and tight agreement between replicates (Figure 3a and Supplementary Table 1).

**Figure 3.**
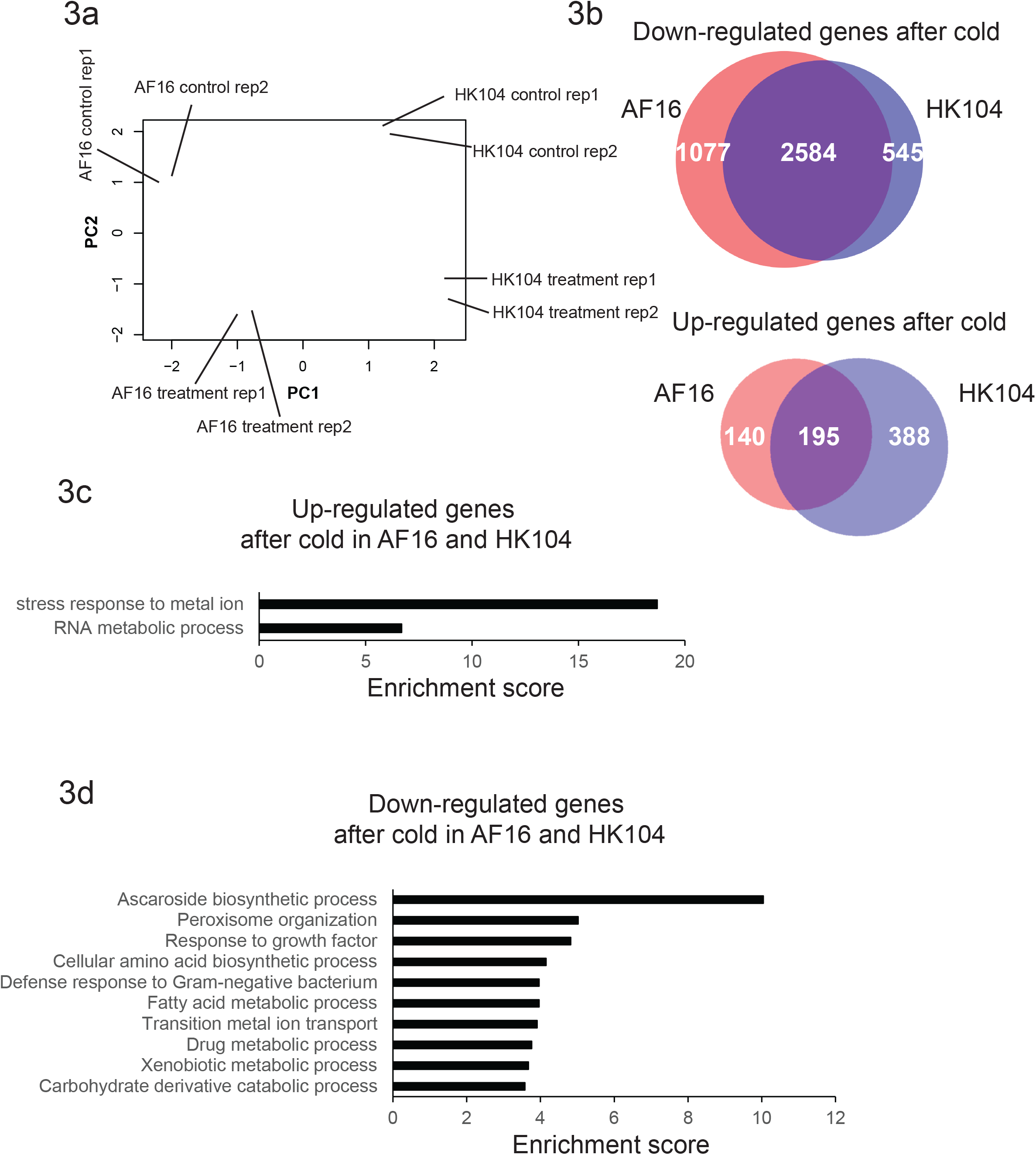
Cold-evoked expression programs shared between AF16 and HK104. **(a)** Shown are the results of principal component analysis of transcriptomes of *C. briggsae* temperate HK104 and tropical AF16 before and after cold treatment (60 hours of incubation at 4°C after development at 20°C). Each point reports values of the first (*x*-axis) and second (*y*-axis) principal component from one replicate of the indicated strain and condition. **(b)** Top, each circle reports the number of genes in the indicated strain with lower expression upon cold treatment than in control conditions. White values report the number of such genes detected only in AF16 (left), only in HK104 (right), or in both strains (center). Bottom, data are as at top except that genes with higher expression in cold conditions were analyzed. **(d)** Each bar reports enrichment (as a ratio of the number observed to the number expected under a genomic null) of genes with the indicated function among those detected in AF16 and HK104 as down-regulated under cold treatment. **(e)** Data are as in **(d)** except that genes with higher expression in cold conditions were analyzed.

We first analyzed AF16 and HK104 separately with respect to cold-evoked changes in gene expression, and inspected commonalities between the strains. In these data, a pattern of declining RNA levels predominated in both AF16 and HK104, with 2584 genes expressed at lower levels upon cold treatment in both strains (at a false discovery rate of 0.15; Figure 3b). Functional-genomic analyses of the latter revealed enrichment for a variety of gene groups involved in growth and cellular metabolism (Figure 3d), as expected if cold-treated animals of both strains slow or arrest many biological processes (Jiang et al., 2018; Robinson & Powell, 2016). By contrast, relatively few genes in each strain were expressed at higher levels in the cold relative to an untreated control (Figure 3b). Only 195 genes exhibited high RNA levels in cold conditions in both AF16 and HK104, with enrichment for functions in RNA processing and metal ion stress (Figure 3c). More salient from these data was the apparent bias for a program unique to temperate HK104, in which we detected an excess of cold-evoked genes that were not called in the analogous tests on AF16 transcriptomes (Figure 3b).

We thus turned our attention to a more rigorous search for strain-by-temperature expression effects (see Methods). The results revealed 191 genes for which the expression response to cold shock was distinct between AF16 and HK104 (at a false discovery rate of 0.15; Supplementary Table 1). These cases of expression divergence were largely specific to cold treatment, with few striking difference between the strains at 20°C. Most (164 genes) followed a pattern of high cold-evoked expression in the temperate strain, HK104, and dropped in expression in the cold in AF16 (Figure 4 and Supplementary Figure 2).

**Figure 4.**
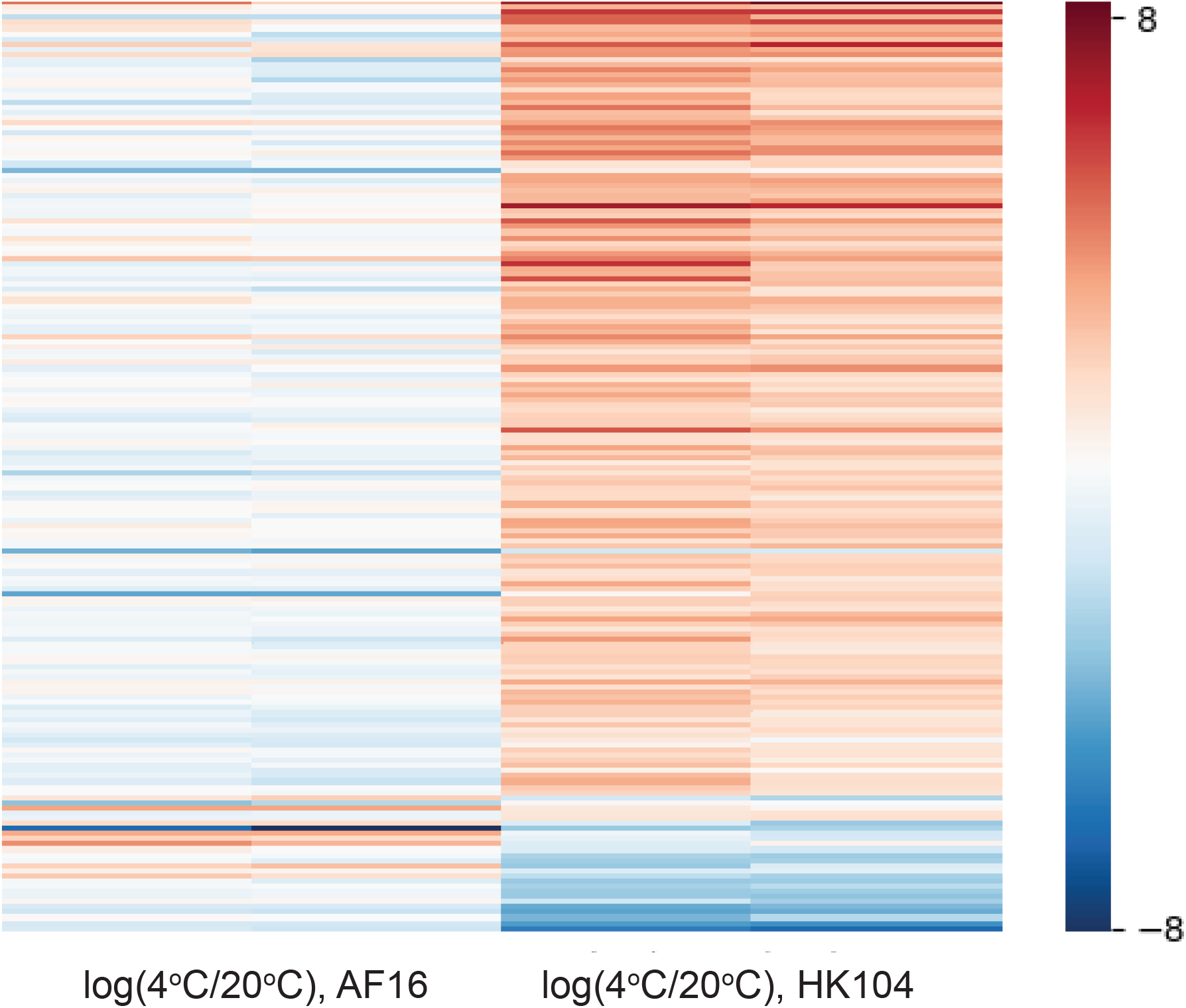
Cold-evoked expression programs that differ between AF16 and HK104. Shown are genes with significant strain-by-temperature effects on expression, in analyses of transcriptomes from **Figure 3**. Each row reports expression measurements for one gene; each column reports a comparison of one replicate of temperate *C. briggsae* HK104 and tropical AF16 treated at the indicated temperature.

We thus formulated a model in which HK104 expressed a unique program of cold-protective factors, whose components could underlie cold resistance at the organismal level. We then earmarked this program for mechanistic follow-up. In transcriptional profiles of AF16 x HK104 hybrid animals, we saw little evidence for *cis*-regulatory divergence at these focal genes (Supplementary Table 1), implicating *trans*-acting variants as the underlying cause of the expression patterns of interest.

### Natural variation and genetics of cold tolerance in *C. elegans*

We set out to use gene ablation to test the phenotypic role of our candidate genes, those expressed at high levels in HK104 during survival of hypothermia. We reasoned that an expedient screen for this purpose could start with *C. elegans.* The latter is much better characterized than *C. briggsae* (Hillier et al., 2007), and genetic mutants are widely available, as opposed to the few mutants generated to date in *C. briggsae* (Hillier et al., 2007); many developmental, behavioral, and physiological phenotypes are conserved between the species (Culp et al., 2020; Hillier et al., 2007; Yin et al., 2018).

To explore the utility of *C. elegans* as a model for cold resistance, we first assayed cold tolerance in 14 wild *C. elegans* strains from temperate locales, and one from a tropical region (Supplementary Table 2). We used a regimen of rearing at 20°C and cold shock at 4°C for 60 hours, in which the *C. elegans* laboratory strain N2 exhibits robust resistance (Ohta et al., 2014). Our results revealed complete lethality in response to cold shock in most other *C. elegans* isolates (Figure 5). Beside N2, originally isolated from England, and CB4555, an N2 descendant (Sterken, Snoek, Kammenga, & Andersen, 2015), only AB1, an Australian isolate previously shown to acclimate rapidly to cold (Okahata et al., 2016), exhibited cold resistance on par with that of temperate-clade *C. briggsae* (Figure 5). We conclude that, in contrast to our observations in *C. briggsae*, temperate collection locality does not associate with cold tolerance in *C. elegans*, strongly suggesting a difference in pressures on the trait between the species. However, we viewed the cold resistance of laboratory *C. elegans* as a useful model for that of temperate *C. briggsae*, with the potential for insights from a genetic screening pipeline.

**Figure 5.**
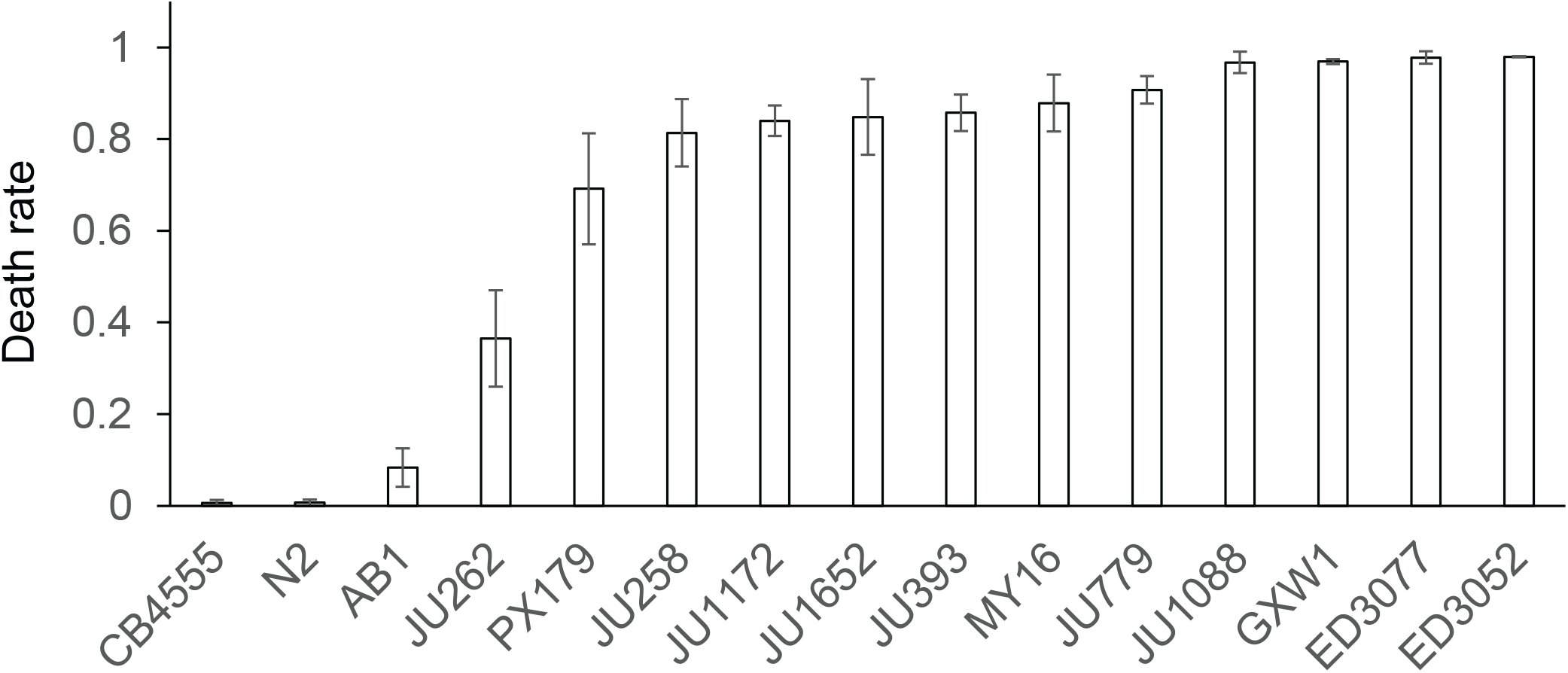
Cold tolerance variation across wild *C. elegans* isolates. Symbols are as in Figure 1 except that wild *C. elegans* strains were analyzed; see Supplementary Table 2 for strain information.

To this end, we carried out a mutant screen in *C. elegans* of cold-induced genes in HK104. To cast the widest possible net for genes of interest, we selected them from our expression data with more lenient cutoffs than we had used for initial genomic analyses (see Methods), amounting to 43 total genes for the screen. For each, we acquired a transgenic strain of the N2 background harboring a mutation in the respective gene, in some cases in a background also including other lesions. Assays for cold resistance revealed seven genes as necessary for the trait in *C. elegans* N2 (Figure 6): *vhp-1* (encoding a MAPK phosphatase), *pgp-8* (an ABC transporter), *ncr-1* (involved in cholesterol trafficking)*, gst-24* (glutathione-S-transferase), *numr-1* (a hypothesized splicing factor), and the uncharacterized genes *M28.8* and *K03H1.5*. By virtue of their role in cold tolerance in *C. elegans*, and their unique cold-evoked induction profile in *C. briggsae* HK104, this set of genes represented our top candidates for determinants of cold resistance in the latter.

**Figure 6.**
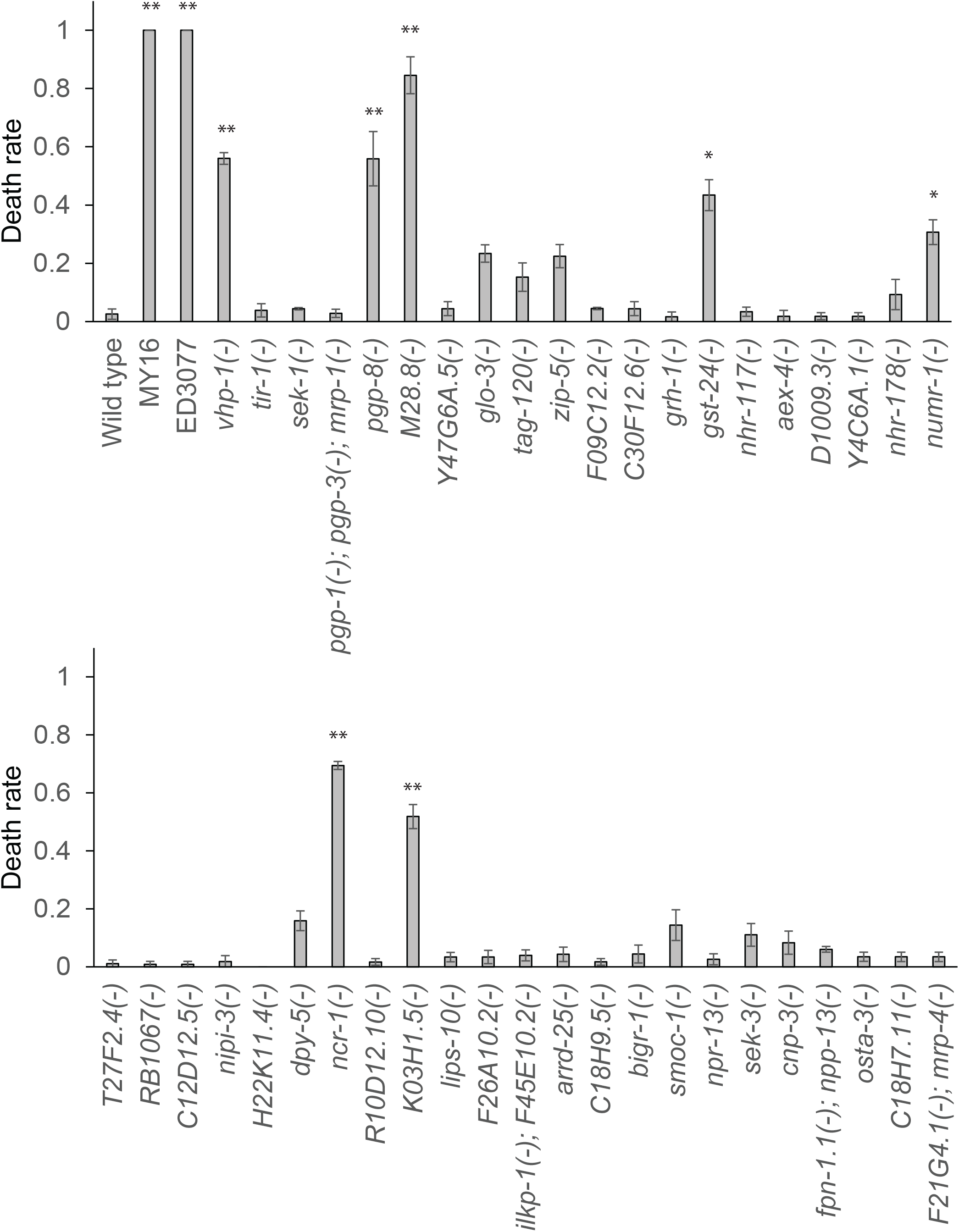
*C. elegans* mutant screen reveals seven genes that impair cold tolerance. Each bar reports cold tolerance in the *C. elegans* laboratory N2 strain (wild-type) or an isogenic strain harboring a mutation (-) in the indicated gene(s). Symbols are as in Figure 1 except that cold treatment was 120 hours. Asterisks report results from a one-sided *t*-test comparing the indicated strain to wild-type N2; *, *p* < 0.05; **, *p* < 0.01.

### Genetic determinants of cold resistance in temperate *C. briggsae*

To explore the phenotypic role of our candidate genes in *C. briggsae*, we made use of a CRISPR-Cas9 system for targeted mutations, for which we chose to focus on five genes with the largest effect size in our *C. elegans* cold resistance screen (*vhp-1*, *pgp-8, M28.8, ncr-1* and *K03H1.5)*. Of these, we were unable to develop homozygous mutants for *ncr-1* in the HK104 temperate *C. briggsae* strain, suggesting an essential function for this gene. For each of the remaining genes in turn (*vhp-1*, *pgp-8, M28.8,* and *K03H1.5*), we established an HK104 line and, separately, a line of tropical *C. briggsae* AF16 harboring a premature stop codon in the coding region (Supplementary Table 3). We then investigated the cold survival phenotype of each such mutant strain. For this purpose, given the extreme cold resistance of HK104 (Figures 1 and 2), we subjected mutants in this background to five days of cold treatment alongside a wild-type control; results revealed a robust and significant increase in cold shock lethality in *pgp-8, M28.8,* and *K03H1.5* mutants, confirming the importance of these genes in the HK104 phenotype (Figure 7a). In the AF16 background, which is radically cold-sensitive (Figures 1 and 2), we were required to use a shorter-duration cold-shock assay design; here we observed no effect of *vhp-1*, *pgp-8, M28.8,* or *K03H1.5* mutation at any timepoint (Figure 7b). These data establish that several of our top genes contribute uniquely to the cold resistance phenotype in HK104, validating our inference from this strain’s expression profiles (Figure 4) as a resource for mechanistic insights in this system.

**Figure 7.**
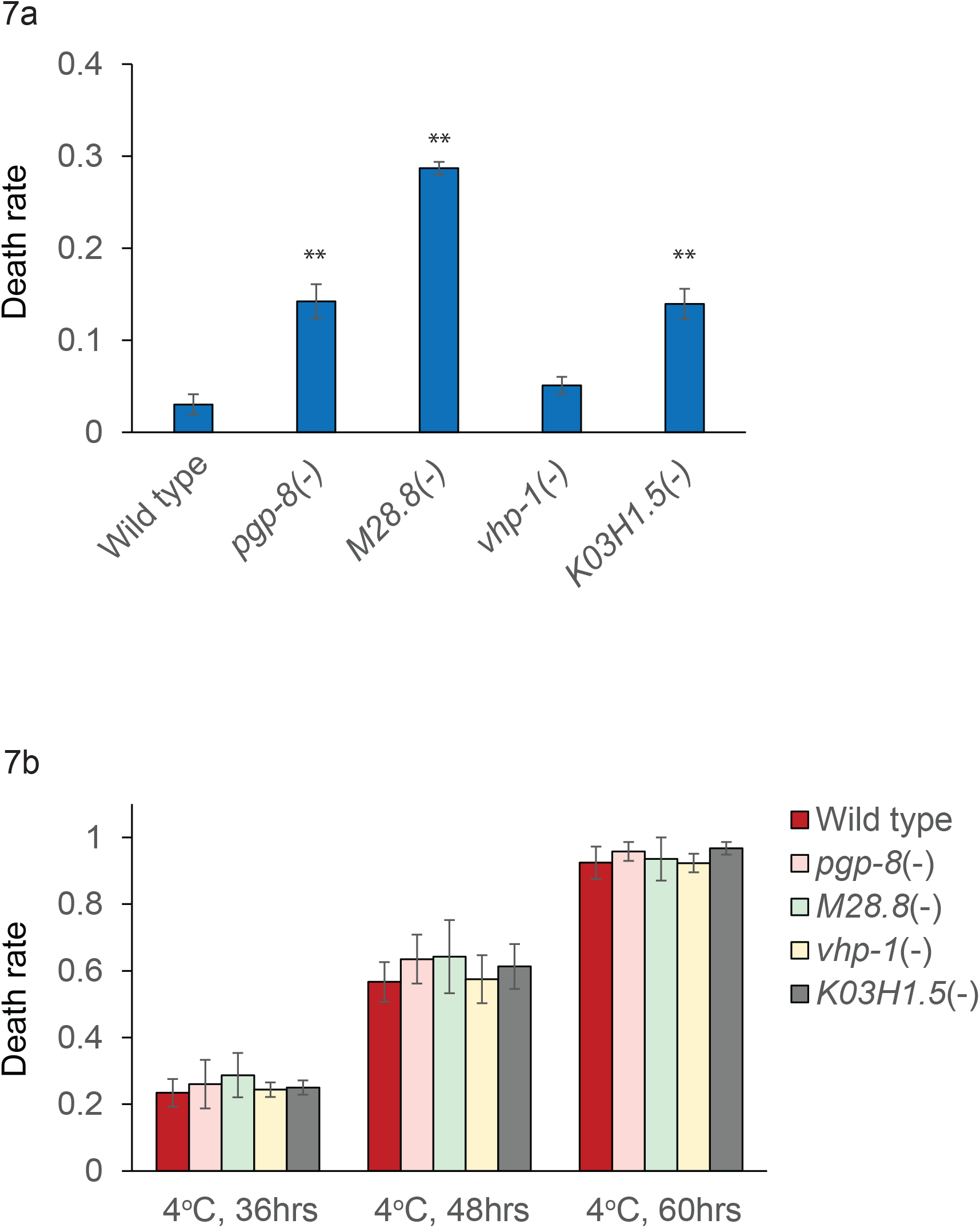
Four genes are important for HK104 cold tolerance. **(a)** Each bar reports cold tolerance in the temperate *C. briggsae* HK104 strain (wild-type) or an isogenic strain harboring a mutation (-) in the indicated gene. Symbols are as in Figure 1 except that cold exposure was for the time indicated. **(b)** Symbols are as in (a) except that the tropical *C. briggsae* AF16 strain was analyzed. Asterisks report results from a one-sided *t*-test comparing the indicated strain to the wild-type of its respective background; *, *p* < 0.05; **, *p* < 0.01. See Supplementary Table 3 for strain information.

## Discussion

Understanding diversity in the natural world requires the discovery of traits that have changed in response to ecological factors, and the use of molecular tools to understand their mechanisms. Heat and cold response are particularly ripe for evolutionary study in ectotherms, whose body temperature depends on that of their environment (Flouris & Piantoni, 2015). In this work, we have established resistance to cold shock as a character distinguishing the temperate clade of the nematode *C. briggsae* from cold-sensitive tropical isolates. Our findings dovetail with the known fecundity disadvantage of temperate *C. briggsae* at high temperature relative to tropical strains (Prasad et al., 2011), as well as behavioral (Cutter et al., 2006; Stegeman et al., 2019, 2013) and transcriptional (Mark et al., 2019) differences between the clades under hot conditions.

Our focus on cold was motivated by the cooler extreme winter temperatures in the collection localities of temperate *C. briggsae* (Adrion et al., 2015; Stegeman et al., 2013) By contrast, the average summer temperature is comparable in most such collection sites, regardless of whether the region is temperate or tropical (Prasad et al., 2011). Given these climatic factors, and the striking phenotype we report here, it is tempting to speculate that selection for cold survival in the winter months has been a key driver of local adaptation in *C. briggsae* as in other nematodes (McGaughran & Sommer, 2014). Any such evolutionary events would have happened remarkably quickly, in light of the 700-year divergence time estimated for the temperate and tropical clades (Cutter et al., 2006).

Evolutionary pressures on the trait in *C. elegans* remain less clear, since we and others (Okahata et al., 2016) have found strains of this species from temperate localities to run the gamut from cold-resistant to cold-sensitive. This may be the product of migration and admixture in the worldwide *C. elegans* population (Petersen, Dirksen, & Schulenburg, 2015), preventing local adaptation in hot or cold environments, or in any one niche (although important exceptions have been reported (Crombie et al., 2019)). As *C. elegans* and *C. briggsae* diverged ~100 million years ago (Hillier et al., 2007), cold tolerance in the two species likely arose independently. The vast divergence in their genomes (Stein et al., 2003) would represent differences in the chassis on which the trait was built, reflected in the fact that some genes we tested affected cold shock survival in resistant *C. elegans* but not *C. briggsae*.

Our evolutionary analysis of wild worms complements an extensive literature on the basic biology of hypothermia response in laboratory *C. elegans.* An important thread of this prior work has highlighted the dieoff of animals re-introduced into warm temperatures after cold shock, and its physiological and transcriptomic correlates (Jiang et al., 2018; Robinson & Powell, 2016). For our expression profiling, instead of the re-warming recovery phase, we focused on the end of the cold exposure regimen, and the genes with high RNA levels at this timepoint. Our validation of a causal role in cold tolerance for seven such genes (in *C. elegans*, temperate *C. briggsae*, or both) suggests that cold resistance hinges in part on physiology during the treatment itself. Under one compelling model, protective factors expressed during cold stress could mitigate tissue damage before recovery even starts. Additionally, proteins expressed in the cold could set up a poised state for rapid signaling and repair during recovery. Ultimately, as a complete picture of the biology of cold resistance becomes clear, it will likely integrate mechanisms operating in the two phases; and the failures in tropical *C. briggsae* could prove to manifest in either one. The transcription factor gene *ZIP-10*, known to promote death during recovery after cold treatment (Jiang et al., 2018), was expressed less in tropical AF16 than in temperate HK104 in our profiles (Supplementary Table 1), suggesting that the trait divergence between the strains hinges on a mechanism distinct from the *ZIP-10* pathway.

That said, among the many other known molecular mechanisms of cold resistance in *C. elegans* (Jiang et al., 2018; Ma et al., 2015; Murray, Hayward, Govan, Gracey, & Cossins, 2007; Okahata, Wei, Ohta, & Kuhara, 2019; Robinson & Powell, 2016; Sonoda, Ohta, Maruo, Ujisawa, & Kuhara, 2016; Takagaki et al., 2020; Ujisawa et al., 2018), some are echoed in the genes we have validated in the trait. For instance, regulating membrane lipid composition and fluidity is well-characterized as a strategy for cold tolerance in the worm (Ma et al., 2015; Murray et al., 2007); the cholesterol processing gene *ncr-1,* which we found to be required for cold tolerance in *C. elegans*, is likely to act through this mechanism. Likewise, detailed studies have established the role of sensory neurons in cold resistance (Ohta et al., 2014; Okahata et al., 2019; Sonoda et al., 2016; Takagaki et al., 2020; Ujisawa et al., 2018). This system could well involve *M28.8*, which we identified as required for the trait in *C. elegans* and temperate *C. briggsae*, is expressed in dopaminergic neurons, and is the ortholog of a *Drosophila* photoreceptor gene (Nie et al., 2012).

Furthermore, in our larger set of cold-evoked expression changes, several emergent patterns parallel those seen in the broader literature. In previous analysis of *C. elegans,* recovery after cold shock was associated with changes in lipid and amino acid metabolism, groups from which many genes exhibited low expression during cold shock itself in our *C. briggsae* study (Jiang et al., 2018). Likewise, RNA binding and stabilization factors are activated by cold in mammalian cells (Sonna, Fujita, Gaffin, & Lilly, 2002) and in yeast and bacteria (Aguilera, Randez-Gil, & Prieto, 2007; Keto-Timonen et al., 2016), just as we have seen in *C. briggsae*. Metal ion stress response, another cold-induced gene set in mammalian cells (Sonna et al., 2002), also featured among upregulated genes in our *C. briggsae* data. Additional stress response genes are induced in cold-shock studies of other organisms, including heat shock factors (Aguilera et al., 2007; Keto-Timonen et al., 2016; Rinehart et al., 2007; Shore et al., 2013; Sonna et al., 2002), whose relevance in the worm remains to be elucidated.

In summary, we have established cold resistance in *C. briggsae* as a rapidly evolved, and likely adaptive, product of ecological diversification, and we have traced the attendant expression changes in genes required for cold tolerance. These findings underscore the power of ecological-genetic studies in wild strains of a tractable model organism, which hold promise for continued progress in the discovery of how nature builds new traits.

## Acknowledgements

This work was supported by NIH GM120430-A1 to RBB and JLG, NIH R35GM119828 to JLG, and NIH 1S10OD021686 to JLG for the COPAS large particle sorter housed in the Buck Institute Morphology and Imaging Core; the American Federation for Aging Research; and the Glenn Foundation for Medical Research. QX1410 and VX34 were kind gifts from Dr. Erik Andersen (Northwestern University). All other strains were provided by the Caenhorhabditis Genome Center, which is funded by the NIH Office of Research Infrastructure Programs (P40 OD010440), and the National BioResource Project of Japan, supported by the Japanese Ministry of Education, Culture, Science, Sports and Technology. We thank members of the Brem laboratory (UC Berkeley) and Garrison laboratory (Buck Institute for Research on Aging) for insightful discussions.

## Data Accessibility

All raw transcriptome data are available at GEO: GSE171725 and SRA: SRP314054.

## Author Contributions

W.W., A.G.F., J.L.G., and R.B.B. designed research; W.W. and A.G.F. performed research; W.W. and A.G.F. analyzed data; W.W., A.G.F., J.L.G., and R.B.B. wrote and edited the manuscript. All authors read and approved the final manuscript.

---

**Supplemental Figure 1.** Symbols are as in Figure 2 of the main text, except that worms were exposed to cold shock for 7 days.

**Supplemental Figure 2.** Shown are genes with significant strain-by-temperature effects on expression, in analyses of transcriptomes from **Figure 3**. Each row reports expression measurements for one gene; each column reports a comparison of one replicate of cold-treated (4°C) and control (20°C) worms of the indicated *C. briggsae* strain.

**Supplemental Table 1. RNA seq analysis.** In the first two tabs, each row reports results from a comparison of expression of the indicated gene in the indicated strain between cold-treated animals and untreated controls. The first column reports log_2_ of the expression fold-change between treatments, as an average over two biological replicates; the remaining columns report, respectively, the F-statistic, nominal *p*-value, and multiple-testing-adjusted *p*-value from the treatment term of a generalized linear model. In the third tab, the first column reports the log_2_ of the expression fold-change between conditions in HK104, divided by the analogous quantity in AF16; the remaining columns report, respectively, the F-statistic, nominal *p*-value, and multiple-testing-adjusted *p*-value from the strain-by-treatment interaction term of a generalized linear model. In the fourth tab, data are analogous to those in the first two tabs except that allele-specific expression in the HK104 x AF16 F1 hybrid was analyzed.

**Supplemental Table 2. Collection localities of wild *C. elegans* strains.**

**Supplemental Table 3.** *C. briggsae* **mutants.** Each pair of rows reports the context of a Cas9-induced mutation (red) in the indicated gene of the indicated *C. briggsae* strain. Uppercase, exonic sequence; lowercase, intronic sequence.

## Notes

### Competing Interest Statement

The authors have declared no competing interest.

## References

Adrion, J. R., Hahn, M. W., & Cooper, B. S. (2015). Revisiting classic clines in Drosophila melanogaster in the age of genomics. Trends in Genetics : TIG, 31(8), 434–444. doi: 10.1016/j.tig.2015.05.006

Aguilera, J., Randez-Gil, F., & Prieto, J. A. (2007). Cold response in Saccharomyces cerevisiae: New functions for old mechanisms. FEMS Microbiology Reviews, 31(3), 327–341. doi: 10.1111/j.1574-6976.2007.00066.x

Baird, S. E., & Stonesifer, R. (2012). Reproductive isolation in Caenorhabditis briggsae: Dysgenic interactions between maternal- and zygotic-effect loci result in a delayed development phenotype. Reproductive Isolation in Caenorhabditis Briggsae: Dysgenic Interactions between Maternal- and Zygotic-Effect Loci Result in a Delayed Development Phenotype, 1(4), 189–195. doi: 10.4161/worm.23535

Crombie, T. A., Zdraljevic, S., Cook, D. E., Tanny, R. E., Brady, S. C., Wang, Y., … Andersen, E. C. (2019). Deep sampling of Hawaiian Caenorhabditis elegans reveals high genetic diversity and admixture with global populations. ELife, 8. doi: 10.7554/eLife.50465

Culp, E., Richman, C., Sharanya, D., Jhaveri, N., Van Den Berg, W., & Gupta, B. P. (2020). Genome editing in the nematode Caenorhabditis briggsae using the CRISPR/Cas9 system. Biology Methods and Protocols, 5(1), 1–5. doi: 10.1093/biomethods/bpaa003

Cutter, A. D., Félix, M. A., Barrière, A., & Charlesworth, D. (2006). Patterns of nucleotide polymorphism distinguish temperate and tropical wild isolates of Caenorhabditis briggsae. Genetics, 173(4), 2021–2031. doi: 10.1534/genetics.106.058651

Flouris, A. D., & Piantoni, C. (2015). Links between thermoregulation and aging in endotherms and ectotherms. Temperature, 2(1), 73–85. doi: 10.4161/23328940.2014.989793

Friedland, A. E., Tzur, Y. B., Esvelt, K. M., Colaiácovo, M. P., Church, G. M., & Calarco, J. A. (2013). Heritable genome editing in C. elegans via a CRISPR-Cas9 system. Nature Methods, 10(8), 741–743. doi: 10.1038/nmeth.2532

Graustein, A., Caspar, J. M., Walters, J. R., & Palopoli, M. F. (2002). Levels of DNA polymorphism vary with mating system in the nematode genus Caenorhabditis. Genetics, 161(1), 99–107. doi: 10.1093/genetics/161.1.99

Hillier, L. D. W., Miller, R. D., Baird, S. E., Chinwalla, A., Fulton, L. A., Koboldt, D. C., & Waterston, R. H. (2007). Comparison of C. elegans and C. briggsae genome sequences reveals extensive conservation of chromosome organization and synteny. PLoS Biology, 5(7), 1603–1616. doi: 10.1371/journal.pbio.0050167

Jiang, W., Wei, Y., Long, Y., Owen, A., Wang, B., Wu, X., … Ma, D. K. (2018). A genetic program mediates cold-warming response and promotes stress-induced phenoptosis in C. elegans. ELife, 7. doi: 10.7554/eLife.35037

Keto-Timonen, R., Hietala, N., Palonen, E., Hakakorpi, A., Lindström, M., & Korkeala, H. (2016). Cold Shock Proteins: A Minireview with Special Emphasis on Csp-family of Enteropathogenic Yersinia. Frontiers in Microbiology, 7(July), 1–7. doi: 10.3389/fmicb.2016.01151

Kraemer, S. A., & Boynton, P. J. (2017). Evidence for microbial local adaptation in nature. Molecular Ecology, 26(7), 1860–1876. doi: 10.1111/mec.13958

Lacher, T. E., & Goldstein, M. I. (1997). Tropical ecotoxicology: Status and needs. Environmental Toxicology and Chemistry, 16(1), 100–111. doi: 10.1897/1551-5028(1997)016<0100:TESAN>2.3.CO;2

Ma, D. K., Li, Z., Lu, A. Y., Sun, F., Chen, S., Rothe, M., … Horvitz, H. R. (2015). Acyl-CoA Dehydrogenase Drives Heat Adaptation by Sequestering Fatty Acids. Cell, 161(5), 1152–1163. doi: 10.1016/j.cell.2015.04.026

Mark, S., Weiss, J., Sharma, E., Liu, T., Wang, W., Claycomb, J. M., & Cutter, A. D. (2019). Genome structure predicts modular transcriptome responses to genetic and environmental conditions. Molecular Ecology, 28(16), 3681–3697. doi: 10.1111/mec.15185

McGaughran, A., & Sommer, R. J. (2014). Natural variation in cold tolerance in the nematode Pristionchus pacificus: The role of genotype and environment. Biology Open, 3(9), 832–838. doi: 10.1242/bio.20148888

Murray, P., Hayward, S. A. L., Govan, G. G., Gracey, A. Y., & Cossins, A. R. (2007). An explicit test of the phospholipid saturation hypothesis of acquired cold tolerance in Caenorhabditis elegans. Proceedings of the National Academy of Sciences of the United States of America, 104(13), 5489–5494. doi: 10.1073/pnas.0609590104

Nie, J., Mahato, S., Mustill, W., Tipping, C., Bhattacharya, S. S., & Zelhof, A. C. (2012). Cross species analysis of Prominin reveals a conserved cellular role in invertebrate and vertebrate photoreceptor cells. Developmental Biology, 371(2), 312–320. doi: 10.1016/j.ydbio.2012.08.024

Ohta, A., Ujisawa, T., Sonoda, S., & Kuhara, A. (2014). Light and pheromone-sensing neurons regulates cold habituation through insulin signalling in Caenorhabditis elegans. Nature Communications, 5, 8–9. doi: 10.1038/ncomms5412

Okahata, M., Ohta, A., Mizutani, H., Minakuchi, Y., Toyoda, A., & Kuhara, A. (2016). Natural variations of cold tolerance and temperature acclimation in Caenorhabditis elegans. Journal of Comparative Physiology B: Biochemical, Systemic, and Environmental Physiology, 186(8), 985–998. doi: 10.1007/s00360-016-1011-3

Okahata, M., Wei, A. D., Ohta, A., & Kuhara, A. (2019). Cold acclimation via the KQT-2 potassium channel is modulated by oxygen in Caenorhabditis elegans. Science Advances, 5(2), 1–13. doi: 10.1126/sciadv.aav3631

Petersen, C., Dirksen, P., & Schulenburg, H. (2015). Why we need more ecology for genetic models such as C. elegans. Trends in Genetics : TIG, 31(3), 120–127. doi: 10.1016/j.tig.2014.12.001

Prasad, A., Croydon-Sugarman, M. J. F., Murray, R. L., & Cutter, A. D. (2011). Temperature-dependent fecundity associates with latitude in caenorhabditis briggsae. Evolution, 65(1), 52–63. doi: 10.1111/j.1558-5646.2010.01110.x

Rinehart, J. P., Li, A., Yocum, G. D., Robich, R. M., Hayward, S. A. L., & Denlinger, D. L. (2007). Up-regulation of heat shock proteins is essential for cold survival during insect diapause. Proceedings of the National Academy of Sciences of the United States of America, 104(27), 11130–11137. doi: 10.1073/pnas.0703538104

Robinson, J. D., & Powell, J. R. (2016). Long-term recovery from acute cold shock in Caenorhabditis elegans. BMC Cell Biology, 17, 2. doi: 10.1186/s12860-015-0079-z

Sanford, E., & Kelly, M. W. (2011). Local adaptation in marine invertebrates. Annual Review of Marine Science, 3, 509–535. doi: 10.1146/annurev-marine-120709-142756

Savolainen, O., Lascoux, M., & Merilä, J. (2013). Ecological genomics of local adaptation. Nature Reviews. Genetics, 14(11), 807–820. doi: 10.1038/nrg3522

Shore, A. M., Karamitri, A., Kemp, P., Speakman, J. R., Graham, N. S., & Lomax, M. A. (2013). Cold-Induced Changes in Gene Expression in Brown Adipose Tissue, White Adipose Tissue and Liver. PLoS ONE, 8(7), 1–9. doi: 10.1371/journal.pone.0068933

Sonna, L. A., Fujita, J., Gaffin, S. L., & Lilly, C. M. (2002). Invited review: Effects of heat and cold stress on mammalian gene expression. Journal of Applied Physiology (Bethesda, Md. : 1985), 92(4), 1725–1742. doi: 10.1152/japplphysiol.01143.2001

Sonoda, S., Ohta, A., Maruo, A., Ujisawa, T., & Kuhara, A. (2016). Sperm Affects Head Sensory Neuron in Temperature Tolerance of Caenorhabditis elegans. Cell Reports, 16(1), 56–65. doi: 10.1016/j.celrep.2016.05.078

Stegeman, G. W., Baird, S. E., Ryu, W. S., & Cutter, A. D. (2019). Genetically distinct behavioral modules underlie natural variation in thermal performance curves. G3: Genes, Genomes, Genetics, 9(7), 2135–2151. doi: 10.1534/g3.119.400043

Stegeman, G. W., De Mesquita, M. B., Ryu, W. S., & Cutter, A. D. (2013). Temperature-dependent behaviours are genetically variable in the nematode Caenorhabditis briggsae. Journal of Experimental Biology, 216(5), 850–858. doi: 10.1242/jeb.075408

Stein, L. D., Bao, Z., Blasiar, D., Blumenthal, T., Brent, M. R., Chen, N., … Waterston, R. H. (2003). The genome sequence of Caenorhabditis briggsae: A platform for comparative genomics. PLoS Biology, 1(2). doi: 10.1371/journal.pbio.0000045

Sterken, M. G., Snoek, L. B., Kammenga, J. E., & Andersen, E. C. (2015). The laboratory domestication of Caenorhabditis elegans. Trends in Genetics : TIG, 31(5), 224–231. doi: 10.1016/j.tig.2015.02.009

Takagaki, N., Ohta, A., Ohnishi, K., Kawanabe, A., Minakuchi, Y., Toyoda, A., … Kuhara, A. (2020). The mechanoreceptor DEG‐1 regulates cold tolerance in Caenorhabditis elegans. EMBO Reports, 21(3), 1–14. doi: 10.15252/embr.201948671

Ujisawa, T., Ohta, A., Ii, T., Minakuchi, Y., Toyoda, A., Ii, M., & Kuhara, A. (2018). Endoribonuclease ENDU-2 regulates multiple traits including cold tolerance via cell autonomous and nonautonomous controls in Caenorhabditis elegans. Proceedings of the National Academy of Sciences of the United States of America, 115(35), 8823–8828. doi: 10.1073/pnas.1808634115

Wang, W., Chaturbedi, A., Wang, M., An, S., Velayudhan, S. S., & Lee, S. S. (2018). SET-9 and SET-26 are H3K4me3 readers and play critical roles in germline development and longevity. ELife, 7, 1–33. doi: 10.7554/eLife.34970

Yin, D., Schwarz, E. M., Thomas, C. G., Felde, R. L., Korf, I. F., Cutter, A. D., … Haag, E. S. (2018). Rapid genome shrinkage in a self-fertile nematode reveals sperm competition proteins. Science (New York, N.Y.), 359(6371), 55–61. doi: 10.1126/science.aao0827

