## Supplemental Info for "Cold survival and its molecular mechanisms in a locally adapted nematode population"

### Figure S1

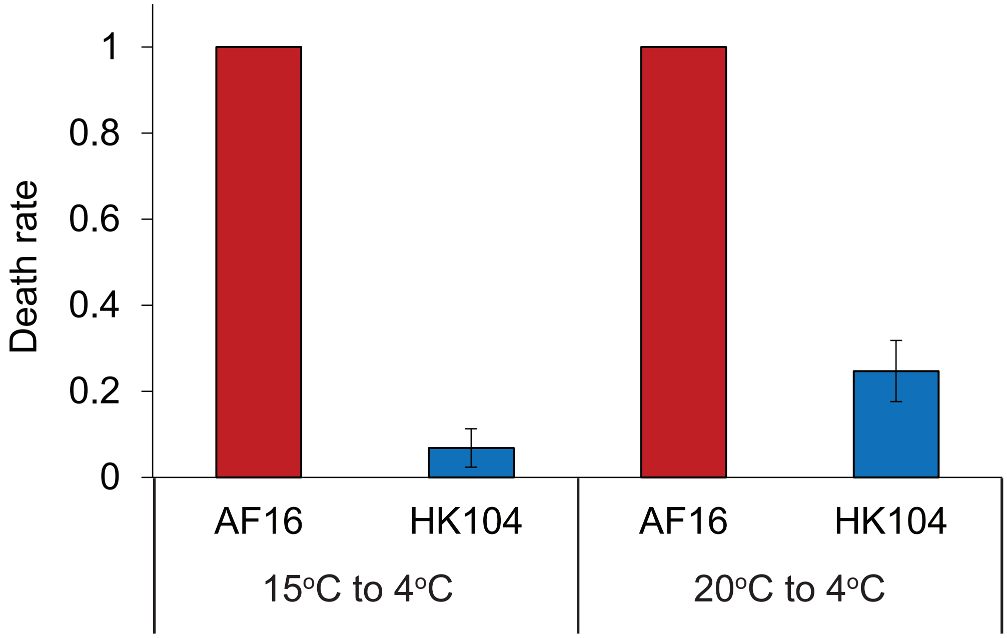

Death rate
1
0.8
0.6
0.4
0.2
0
AF16
HK104
AF16
HK104
15oC to 4oC
20oC to 4oC

### Figure S2

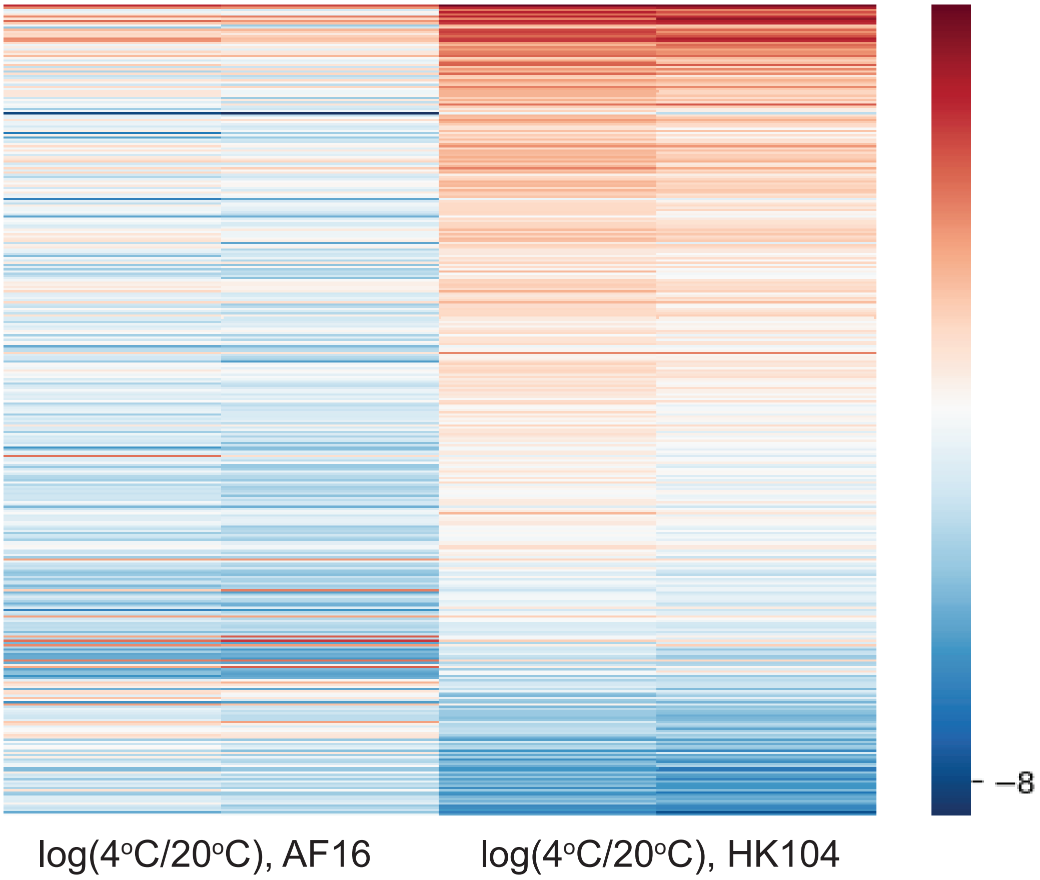

### Supplemental Table S2

| Strain | Locality of origin | Latitude/longitude | Climate |
|---|---|---|---|
| CB4555 | Pasadena, Ca | 34.1478° N, 118.1445° W | Temperate |
| N2 | Bristol, England | 51.4545° N, 2.5879° W | Temperate |
| AB1 | Adelaide, Australia | 34.9285° S, 138.6007° E | Temperate |
| JU262 | Indre, France | 46.6614° N, 1.4483° E | Temperate |
| PX179 | Eugene, OR | 44.0521° N, 123.0868° W | Temperate |
| JU258 | Madeira, Ribeiro Frio | 32.7333° N, 16.8833° W | Temperate |
| JU1172 | Concepcion, Chile | 36.8201° S, 73.0444° W | Temperate |
| JU393 | Hermanville, France | 49.2865° N, 0.3138° W | Temperate |
| JU1652 | Montevideo, Uruguay | 34.9011° S, 56.1645° W | Temperate |
| MY16 | Munster, Northwest Germany | 51.9607° N, 7.6261° E | Temperate |
| JU779 | Lisbon, Portugal | 38.7223° N, 9.1393° W | Temperate |
| GXW1 | Hubei Province, China | 30.7378° N, 112.2384° E | Temperate |
| ED3077 | Nairobi, Kenya | 1.2921° S, 36.8219° E | Tropical |
| ED3052 | Ceres, South Africa | 33.4007° S, 19.2950° E | Temperate |
| JU1088 | Shizuoka prefecture, Japan | 35.0929° N, 138.3190° E | Temperate |

### Supplemental Table S3

| | | |
|---|---|---|
| AF16 wild type (WBGene00025434) | TCACGCTCTCTTCATGCGGGTTGGCGGCACTCATTCGTGAGGCTCCA | -7bp |
| AF16 vhp-1(-) (WBGene00025434) | TCACGCTCTCTTCATGCGGGCACTCATTCGTGAGGCTCCA | |
| HK104 wild type (WBGene00025434) | ACACAGTCACGCTCTCTTCATGCGGGTTGGCGGCACTCATTCGTGAG | -7bp |
| HK104 vhp-1(-) (WBGene00025434) | ACACAGTCACGCTCTCTTCATGGCGGCACTCATTCGTGAG | |
| AF16 wild type (WBGene00037162) | GAGAGAGGTACTCAACTTTCTGGCGGTCAAAAACAACGAA | +8bp |
| AF16 pgp-8(-) (WBGene00037162) | GAGAGAGGTACTCAACTTTCGGTCAACTTGGCGGTCAAAAACAACGAA | |
| HK104 wild type (WBGene00037162) | GAATTGCAATTGCTCGTGTTTTGGTGAAAAACCCGAAAATCCTTCTTTTG[illegible] | -491bp |
| HK104 pgp-8(-) (WBGene00037162) | GAATTGCAATTGCTCGTGTTTGGCAGATTCTCAGAGAATG | |
| AF16 wild type (WBGene00025987) | TCCCAACAATCAACACCTCCACCTGAGCAATGTGATTTCGACTCACCAGC[illegible] | -281bp |
| AF16 M28.8(-) (WBGene00025987) | TCCCAACAATCAACACCTCCGGTGGAGCTGTTACCTATGA | |
| HK104 wild type (WBGene00025987) | GTCCCAACAATCAACACCTCCACCTGAGCAATGTGATTTCGA | +1bp |
| HK104 M28.8(-) (WBGene00025987) | GTCCCAACAATCAACACCTCAATCCTGAGCAATGTGATTTCGA | |
| AF16 wild type (WBGene00031437) | CCGCCCTTTTCTGCTCCTCCATGGACGTTTCCAAATCCATCATGGCCGAAA[illegible] | -54bp |
| AF16 K03H1.5(-) (WBGene00031437) | CCGCCCTTTTCTGCTCCTCCgtatttcctctagtagaagc | |
| HK104 wild type (WBGene00031437) | CGCCCTTTTCTGCTCCTCCATGGACGTTTCCAAATCCATC | +5bp |
| HK104 K03H1.5(-) (WBGene00031437) | CGCCCTTTTCTGCTCCTCCAGCAGAACGACGTTTCCAAATCCATC | |
